# The anatomical distribution of genetic associations

**DOI:** 10.1101/021824

**Authors:** Alan Wells, Nathan Kopp, Xiaoxiao Xu, David R. O’Brien, Wei Yang, Arye Nehorai, Tracy L. Adair-Kirk, Raphael Kopan, J.D. Dougherty

**Author notes:** equal contribution of authors (Alan Wells, Nathan Kopp). Contact: Dr. Joseph Dougherty Department of Genetics Campus Box 8232 4566 Scott Ave. St. Louis, MO. 63110-1093 E.

## Abstract

Deeper understanding of the anatomical intermediaries for disease and other complex genetic traits is essential to understanding mechanisms and developing new interventions. Existing ontology tools provide functional annotations for many genes in the genome and they are widely used to develop mechanistic hypotheses based on genetic and transcriptomic data. Yet, information about where a set of genes is expressed may be equally useful in interpreting results and forming novel mechanistic hypotheses for a trait. Therefore, we developed a framework for statistically testing the relationship between gene expression across the body and sets of candidate genes from across the genome. We validated this tool and tested its utility on three applications. First, using thousands of loci identified by GWA studies, our framework identifies the number of disease-associated genes that have enriched expression in the disease-affected tissue. Second, we experimentally confirmed an underappreciated prediction highlighted by our tool: variation in skin expressed genes are a major quantitative genetic modulator of white blood cell count - a trait considered to be a feature of the immune system. Finally, using gene lists derived from sequencing data, we show that human genes under constrained selective pressure are disproportionately expressed in nervous system tissues.

## Introduction

Gene annotation tools, such as Gene Ontologies(GO)^1^ and the KEGG database, have long provided an essential foundation for a large number of data mining and pathway analysis tools^2–5^. However, these resources, largely do not incorporate systematic information about gene expression. Every cell in the human body harbors a nearly identical copy of the genome, yet each tissue only expresses the subset of genes that are required to fulfill its particular functions. Thus, knowledge of where a gene is expressed often provides insight into potential functional roles. Furthermore, human diseases are often quite clearly caused by dysfunction or disruption of particular tissues, but human disease genetics typically treat all regions of the genome equally in regard to their prior probability of contributing to disease risk. If one could establish the extent to which genes with highly enriched or specific expression in a particular tissue contribute to diseases of that tissue, then tissue specific expression might also be used to improve our ability to identify risk loci. More immediately, such a method could provide insight to the physiology of poorly understood traits and diseases as well as elucidate new and interesting relationships between our traits and our anatomy. Therefore, we set out to 1) formally test this ‘selective expression’ assumption/hypothesis: that disease genes are over-represented by enriched expression in the disease-relevant tissue, and 2) to establish a foundation, analogous to Gene Ontologies, for tools that could be used to identify potentially interesting novel relationships between human traits and tissues.

A rigorous test of the selective expression hypothesis requires 1) genome wide analysis of gene expression across a large number of tissues, 2) application and validation of an analytical method to identify genes enriched in each tissue, and 3), a comparison to expression data of a large number of trait-associated genes across a large number of traits and disorders.

For a gene expression data set, we leveraged the availability of the Genotype-Tissue Expression (GTEx) project, which includes a previously completed analysis of 1,839 RNAseq samples spanning 45 tissues across 189 post-mortem individuals ^6^. For an analytical method, we adapted our previously developed specificity index statistic ^7^. This statistic was originally designed to identify transcripts enriched in a particular cell type using a large set of cell-specific microarray data profiling a range of cell types in the mouse brain ^8^, and was validated with extensive independent analysis of *in situ* hybridization data ^7^. We recently used this microarray data to design a tool (Cell-type Specific Expression Analysis-CSEA) to test the selective expression hypothesis specifically for cell types of the brain and neurogenetic disease ^9^. Here, we adapt this statistic for use with the GTEx data, validate these tissue specific enrichment lists with Gene Ontologies analysis, and develop an analogous tool (Tissue Specific Expression Analysis – TSEA) for testing the selective expression hypothesis across the body. Finally, we used the TSEA framework to test the selective expression hypothesis using the results of over 100 annotated GWA studies, from the curated NHGRI GWAS catalog ^10^.

Using the expression pattern of trait-associated genes, our tool successfully identifies the tissues for many genetic diseases and complex traits with known anatomical substrates, overall assigning 57 of the traits to one or more tissues, typically with high face validity (Movie S1). Randomization tests reveal that this ability to map particular traits to tissues occurs far more frequently than expected by chance, providing clear statistical support for the selective expression hypothesis overall, and providing a path towards the estimation of appropriate biological priors for future reweighting candidate disease genes by tissue specific expression. In addition, our tool identified a relationship between human skin gene expression and white blood cell count, suggesting that skin integrity is a major modifier of white blood cell count in humans. This hypothesis is inline with previous work in mice showing skin-specific disruption of notch signaling or stratum corneum formation can alter lymphoblast ^11^ and granulocyte ^12^ proliferation. Here, we show that white blood cell count is strongly predicted by inside-out skin barrier function across a range of genetic mutations in distinct molecular pathways. This proof-of-principle analysis shows the utility of our method in helping to rapidly identify the relevant tissues and highlight anatomical relationships for particular traits from human genetic data. Finally, to demonstrate our approach can be broadly useful for interpreting gene lists derived from any data sources beyond GWAS, we show that human genes under constrained selective pressure are disproportionately expressed in nervous system tissues.

## Results

### Identification of transcripts enriched in each tissue

As a source of data to identify selectively expressed and enriched transcripts across tissues, we leveraged a publically available analysis of the GTEx RNAseq data. The GTEx project has assessed the expression of 18,056 protein-coding genes across 45 tissue types. Some of these tissues are different dissections of a larger tissue (e.g., the substantia nigra and frontal cortex are regions of the brain). Still other tissue samples come from different anatomical branches of a larger tissue system (e.g. both aorta and capillaries are blood vessels). While the data are derived from numerous individuals, comparing the observed summarized within tissue variance across all genes, to a permuted set of gene expression, it is evident that variation in gene expression is largely driven by differences between tissues (Figure S1). In other words, the gene expression in one individual’s liver is far more similar to the gene expression in a second liver than it is to gene expression in that individual’s brain. Thus, we used all available data to summarize gene expression in each tissue to a single mean measurement across all individuals to provide a reasonable estimate of the normal expression of this gene in each tissue across the population.

We next used the previously validated pSI algorithm^7^ to identify transcripts enriched in each tissue (Figure S2). In brief, each tissue, was compared, pairwise, to all other available tissues and a ratio was calculated for the expression of every gene. For each comparison, the ratios are converted to ranks to normalize the range, and this rank is averaged across all comparisons for a given tissue, yielding the Specificity Index (SI) – the average rank of the gene on every pairwise tissue comparison list. SI can be assigned a p-value (pSI) using a permutation-based strategy. This algorithm serves to both order the genes from the most highly enriched in a particular tissue to more ubiquitous, and provides the ability to choose particular thresholds to generate lists of genes with enriched expression in each tissue at varying statistical stringencies. Here we use four pSI thresholds (0.05, 0.01, 0.001, 0.0001). The algorithm performed as expected as more stringent thresholds produced smaller yet more specific gene lists, as illustrated with a heatmap for counts of the genes that are shared between each pairwise tissue comparison (Figure S2).

Though the pSI algorithm is relatively robust to minor changes in the composition of the inputted dataset ^9^ we were concerned that including multiple sub-tissue dissections may slightly detract from our ability to identify tissue-enriched genes more broadly. Therefore we focused on a ‘whole-tissue’ version of the analysis where all brain dissections (and similar sub-dissections of other organs) were averaged to a single measure prior to calculation of pSI. This served to slightly increase the number of genes detected as enriched in the whole-tissues at a given pSI threshold and simplified future interpretation to only 25 tissues for the remainder of the analyses. Consistent with earlier reports using microarray data from a smaller number of tissues ^13^ by far the most unique tissue in either analysis was the testis, with 560 highly enriched (pSI<0.0001) transcripts, with the second most unique tissue being the brain (193 transcripts at the same threshold).

Overall, there were 6,922 transcripts detected as modestly enriched (pSI<0.01) in any tissue, with high expression in one tissue and some expression in related tissues (Figure 1A). For example, the brain, the adrenal gland, and the pituitary all release synaptic vesicles full of chemical transmitters, and all share in their enrichment for a number of transcripts related to this process (e.g., *CPLX2*: brain pSI<1E-6, adrenal gland: pSI<0.02; pituitary: pSI<0.002). On average, a transcript showing a pSI<0.01 in at least one tissue might also show a modest enrichment (pSI<0.1) in 2.6 other tissues (+/ – 1.6 s.d.). Using a more stringent threshold (pSI<0.0001) identifies a smaller number of transcripts in all (1,301 across all tissues), but much more likely to have specific expression in a single tissue (Figure 1B and S2). 564 of these highly tissue enriched transcripts don’t even show a trend (pSI<0.1) towards enrichment in any other tissue. These transcripts with highly enriched expression in one tissue include clear examples known to be essential to the functioning of particular tissues (e.g., Figure 1C), a finding supported by a Gene Ontological (GO) analysis. For example, using the DAVID tool^4^, the 501 genes with pSI <0.001 in brain are massively enriched for a large number of biological process terms related to the functioning of the nervous system (e.g., Synaptic Transmission, p-value<2.6E^-56^, B-H corrected; Neurological Systems Process, p-value<2.4E^-33^), while the 1,132 genes with pSI<0.001 from the testis are enriched for the GO terms like Sexual Reproduction(p-value<1E^-64^), or Spermatogenesis(p-value<1E^-56^). Thus, overall the algorithm seemed to perform well in selecting gene sets enriched in each tissue. Nonetheless, to make sure our gene lists were robust to the particular algorithm used we also tested a second method to determine specificity, a variation of Shannon entropy^14^. Overall, the SI and entropy values for gene expression within a tissue are highly correlated (Figure S3). However, the transcript lists determined by the Shannon entropy are smaller subsets of the transcript lists produced by the pSI method at the same threshold, so we utilized the pSI for the remaining analyses.

**Figure 1.**
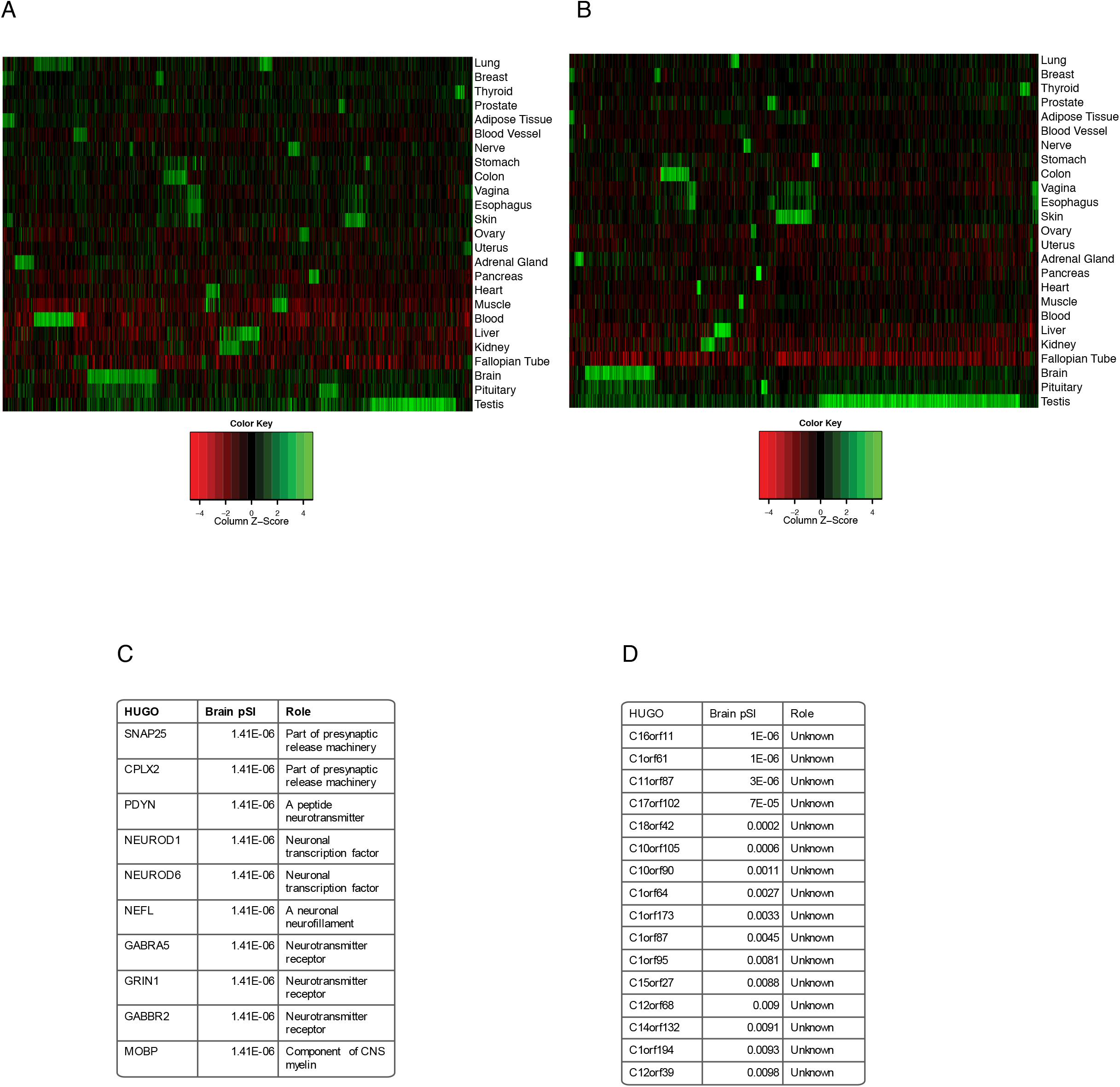
The pSI algorithm can be used to identify transcripts enriched in each tissue. A) Clustering of genes modestly enriched (pSI<0.01) in any tissue, reveals some tissues are more unique (e.g., testis, blood, brain), though some transcripts are found across tissues with related functions. B) Transcripts identified as highly enriched (pSI<0.0001) by pSI clearly show high expression in fewer tissues. C) Examples of annotated genes identified by pSI as highly enriched in the brain include many known neuronal genes. D) Examples of unannotated genes that also show specificity and enrichment of brain expression.

Note that relative to GO, our analysis also provides information for poorly annotated and largely unstudied transcripts (Figure 1D), which won’t be represented in the GO databases. For these, a variation of ‘guilt-by-association’ logic (guilt-by-expression) would suggest that they might also be important for the particular tissue in which they are enriched. To facilitate future investigations of these transcripts from any tissue, we provide a matrix of pSI values for all tissues and genes as Table S1.

### Development and validation of a tissue specific enrichment analysis tool (TSEA)

We next developed a tool to perform an enrichment analysis using the tissue specific expression information. The purpose of this tool is to identify whether a set of ‘candidate genes’ for a trait of interest are disproportionately transcribed in a particular tissue. This is meant to be analogous to existing tools for identifying overabundances of particular GO or KEGG terms in gene lists ^2–5^, and implements a similar statistical framework: Fisher’s Exact test ^15^ coupled with Benjamini-Hochberg (B-H) multiple testing correction ^16^. However, there are two unique properties of our TSEA vis-à-vis GO enrichment. First, unlike GO terms, expression data is not built upon manual curation – thus even poorly annotated and unstudied genes, such as those in Figure 1D, have the opportunity to contribute signal to a TSEA analysis. Second, the pSI statistic provides for a nuanced definition of ‘tissue-specific’ or ‘tissue-enriched’ transcripts. This is important because, while we are testing for an enrichment, *a priori* there is no clear expectation of exactly how uniquely disease genes ought to be expressed in the tissue of interest. For example, mutations in the cystic fibrosis transmembrane conductance regulator gene *CFTR,* cause cystic fibrosis, which effects many different tissues including the lungs, pancreas, and kidney. Though not widely expressed, setting too stringent an expression threshold might exclude this gene from one or the other relevant tissue specific lists (pSI kidney=0.042, lung=0.027, pancreas=0.0011, colon=0.003). Thus, currently the tool is set up to vary pSI systematically and report Fisher’s exact test results at multiple pSI thresholds. Graphically, these results are displayed as a set of concentric hexagons for each tissue, with each smaller hexagon representing a more stringent pSI threshold (Figure 2A). The size of the hexagon is scaled to the number of genes meeting the pSI threshold, and its color indicates results of the Fisher’s exact test. Figure S4 offers a detailed description of how to interpret these plots. For example, we can query the tool with the 14 genes annotated by GO as functioning in Renal Water Homeostasis (GO:0003091), or the 66 genes associated with T Cell Activation (GO:0002286), and rapidly identify that these genes are enriched in the Kidney (Figure 2B) and Blood (Figure 2C), respectively. Note that GO Immune-related terms often map via TSEA to the blood and the lung, both tissues in which lymphoblasts are expected to contribute a substantial portion of the mRNA. Likewise, transcripts for GO terms like glycogen metabolic process are seen to be enriched in multiple tissues known to have high levels of glycogen metabolism (Figure S4).

**Figure 2.**
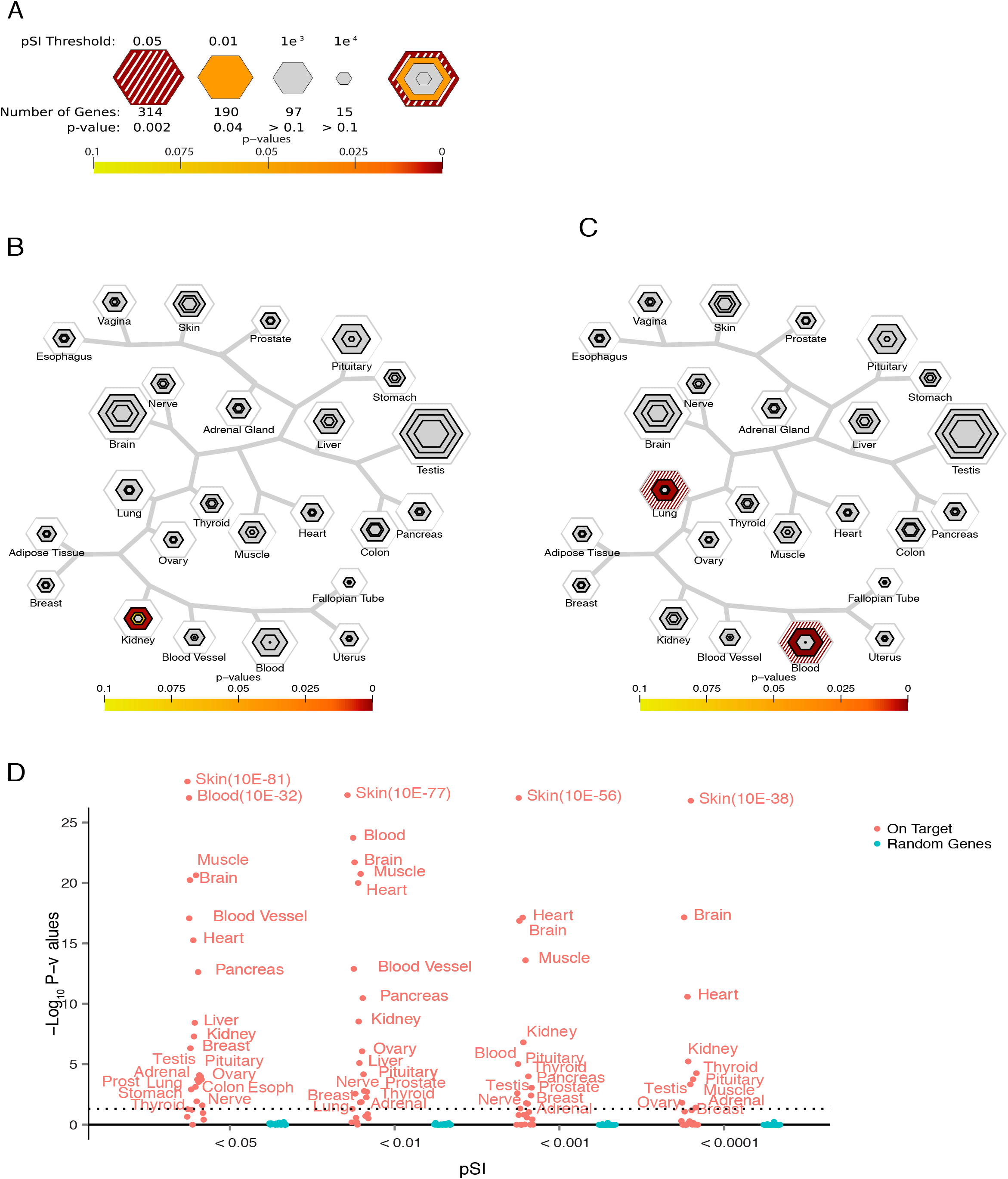
Tissue specific expression analysis (TSEA) correctly identifies tissues related to known biological processes. A) Key for TSEA. Largest hexagon represents pSI<0.05 gene list, and is partially shaded with white to indicate the relative permissiveness of this criterion. More stringent subsets of transcripts enriched in the same tissue are represented by smaller embedded hexagons. All hexagons are scaled to the number of transcripts at that pSI value, and color-coded by p-value of overlap with inputted TSEA ‘candidate gene’ list. B) TSEA using the Gene Ontology (GO) term ‘Renal Water Homeostasis’ as the candidate list indicates that these transcripts have enriched expression in the kidney. C) TSEA for GO term ‘T cell activation involved in immune response’ reveals enrichment for these transcripts in the blood and the lungs, tissues with a substantial proportion of lymphoblasts. D) Across TSEA for GO terms for the development of each tissue, 20/25 GO term gene lists were correctly ascribed to the relevant tissue by TSEA (orange dots, p-values in – log10 scale), while 0/25 same-sized random lists of genes identified any tissue by TSEA at any pSI threshold (blue dots). Horizontal dotted line represents – log10(0.05).

To more thoroughly validate both the functioning of the tool and the pSI statistic, we conducted a systematic analysis of gene lists derived from the GO resource. As GO does not currently include expression information, or a broad term for the genes used in the functioning of a particular tissue in the adult, our best choice as a ‘positive control’ gene list for each tissue was the set of genes annotated as being involved in the development of that particular tissue. For example, GO term 0060537, Muscle Development, is associated with 320 human genes. Though the GTEx data is based on adult tissue, a large subset of these genes is clearly still expressed after development: there are robust statistical signals by TSEA in the muscle at all pSI thresholds (B-H corrected Fishers Exact test p-values of <0.0005 to 10E^-21^). Across GO terms for all tissues the relevant tissue was correctly identified by TSEA in at least one pSI threshold in 20 out of 25 tissues. The remaining tissues each had fewer than 30 genes associated with their GO terms. Compared to this, 0 out of 25 random lists of genes identified any tissue by TSEA at any threshold (Figure 2D). Thus, via the adult GTEx expression information alone, the TSEA tool can accurately identify tissues from a list of genes relevant to their development. This test validates both the functioning of the tool and the quality of the pSI derived lists of tissue specific genes, and suggests the tool will be equally useful at identifying tissue-enriched expression from lists of candidate disease genes, providing an opportunity to test the long-held selective expression hypothesis.

### Test of the selective expression hypothesis

To test the selective expression hypothesis, we applied TSEA to every trait in the publically available GWAS catalog with a reasonable number of genes reported by the authors of the study (>30 genes, a threshold suggested by prior power modeling analyses^9^). Using these 98 ‘candidate gene’ lists, the results from the TSEA are consistent with the known biology for many of the complex and disease associated traits (Figure 3 & Figure 4). Genes associated with metabolic traits map to their tissues of origin. For example, urate levels ^17–25^ associated genes map to the kidney and bilirubin, a blood metabolite indicative of liver function, maps to the liver (Figure 3A & B). Similarly physical traits, such as heart rate ^26–32^, map to the responsible tissue (Figure 3C). Furthermore, in agreement with the recent observation that obesity genes include many genes with high transcription in the brain ^33^, we see clear enrichment of expression in the brain for genes controlling BMI ^34,35^ (Figure 3D). Most autoimmune diseases, such as inflammatory bowel disease ^36–41^, and rheumatoid arthritis ^42–54^ map strongly to tissues containing a high proportion of immune cells (blood and lungs, Figure 4A & B), with some occasional signal in the tissue targeted by auto-immune attack. Chronic kidney disease shows signal in the kidney as well as the thyroid, liver, and prostate, highlighting the affected tissue as well as possibly deranged tissues affecting the pathogenesis of the disease (Figure 4C). Finally, genes associated with cognitive decline in Alzheimer’s disease ^55^ are disproportionately expressed in the brain (Figure 4D), as are other genes associated with psychiatric disease and cognitive function (Movie S1). There is also some suggestive signal in the pituitary, another tissue that undergoes substantial synaptic release, though it is not clear whether this is because it is independently involved or that it simply shares some similarity of gene expression with neurons (Figure 1). TSEA results for any significant trait across the 98 candidate gene lists are found in Movie S1.

**Figure 3.**
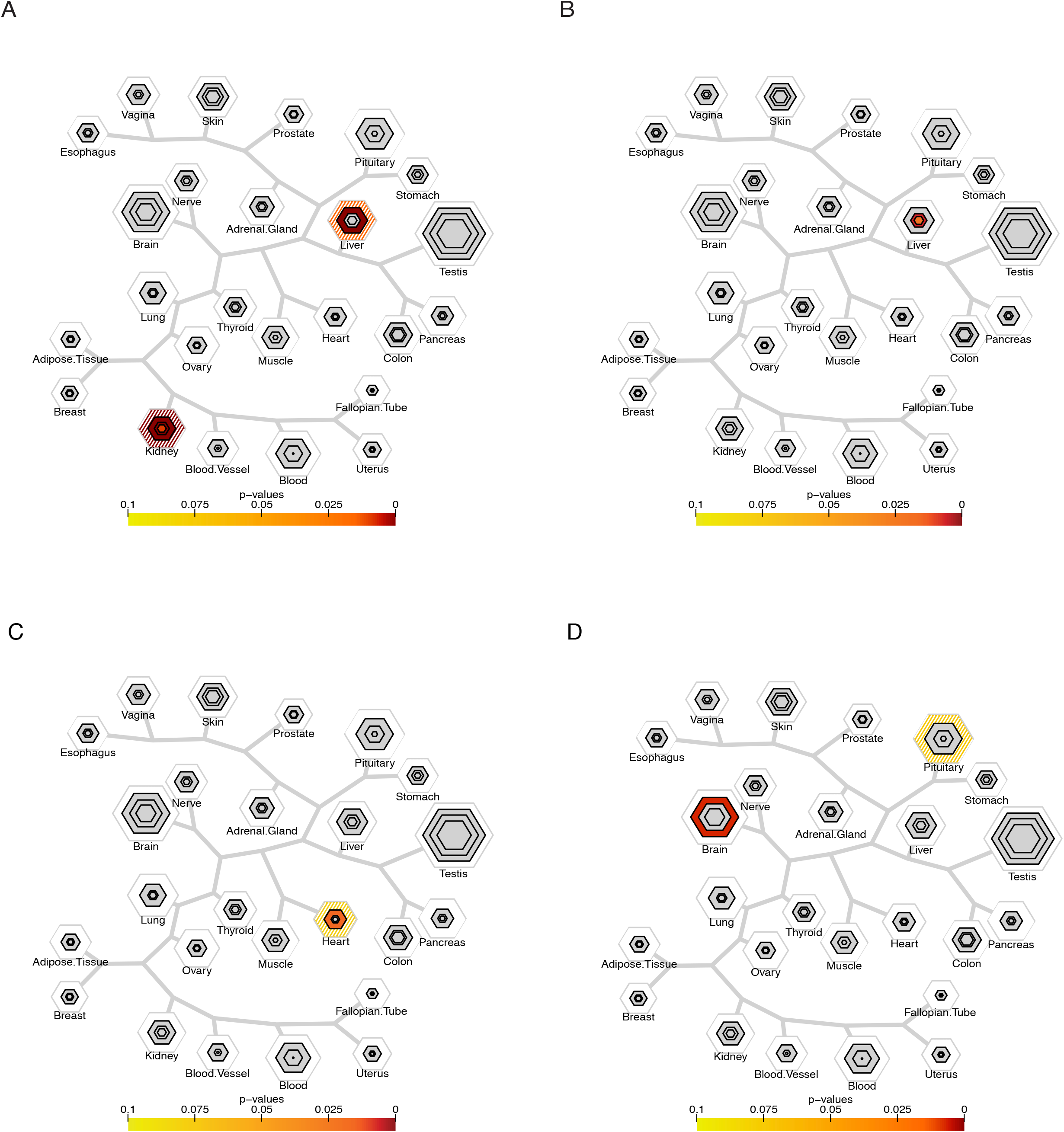
Tissue specific expression analysis identifies tissues associated with human complex traits. A) Genes associated with urate acid levels are disproportionately transcribed in the kidney, with signal in the liver as well. B) Genes associated with Bilirubin levels are disproportionately enriched in the liver. C) Genes regulating heart rate are disproportionately enriched in the heart. D) Genes regulating Body Mass Index are disproportionately transcribed in the brain, with some suggestive signal in the pituitary.

**Figure 4.**
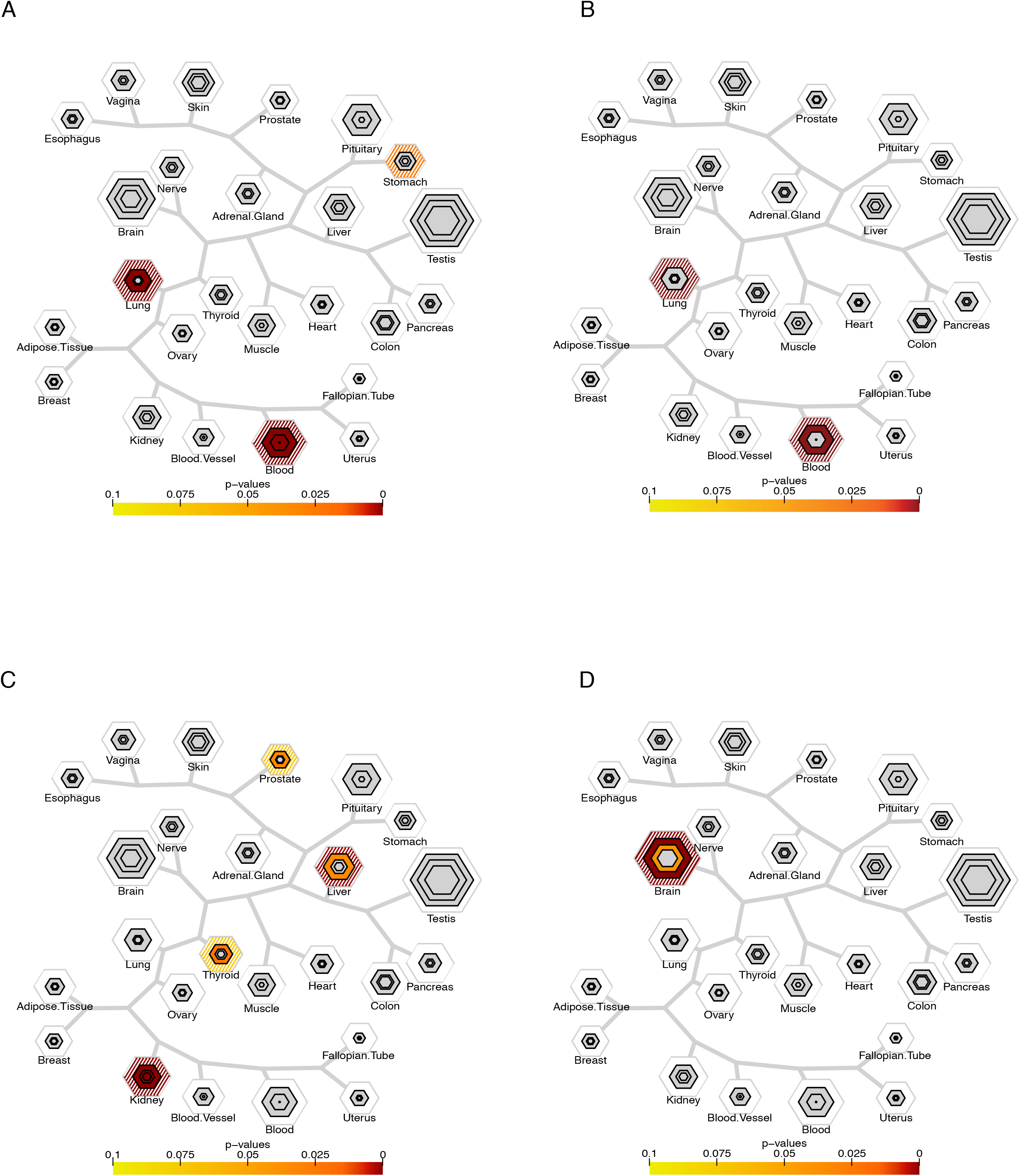
Tissue specific expression analysis identifies tissues associated with human disease traits. A) Autoimmune diseases, such as IBD, show enrichment for transcripts found in blood and lung. There is also suggestive signal in stomach. B) Rheumatoid arthritis shows an overrepresentation of transcripts expressed in the blood and lung. C) Genes associated with chronic kidney disease show signal in the kidney, thryoid, prostate, and D) Genes associated with cognitive decline in Alzheimer’s are disproportionately transcribed in the brain.

Overall, our analyses identified significant overlap between candidate gene lists and transcript-enriched lists for a tissue for 57 out of the 98 traits, providing general support for the selective expression hypothesis across a large number of traits. The tissues that were found significant for each trait are often consistent with the known biology of the trait

To determine if the TSEA method is detecting a larger number of tissue specific relationships than expected by chance (a prediction of the selective expression hypothesis), we repeated the TSEA analysis on 98 more ‘traits’ with randomly constructed candidate gene lists and repeated the process 1000 times. The distribution in Figure 5A shows the number of randomly constructed ‘traits’ with a significant overlap with at least one tissue. The maximum number of significant hits from the thousand random trials is 21 out of 98 traits for the whole-tissue dataset (median:8/98). Therefore the ‘true’ set of 57/98 GWAS traits are mapping to specific tissues far more than expected by chance (p-value<0.001). Furthermore, we compared the distribution of the – log_10_(p-values) from the randomly constructed ‘traits’ and the true GWAS traits (Figure 5B). The – log_10_(p-values) generated by the random gene lists have a small range (min = 1.30, max = 5.55,median=1.58) and tend toward lesser significance. In contrast, the GWAS candidate gene lists produce -log_10_(p-values) that have a wider range (min=1, max=27.07, median=2.57) and a longer tail towards greater significance. A Wilcoxon Rank Sum test supports that the two distributions are different (p < 2.2E-16). Similar results are seen when using the sub-tissue dataset (data not shown). This firmly supports the hypothesis, across a diverse range of phenotypes, that trait associated genes have enriched expression in particular tissues.

**Figure 5.**
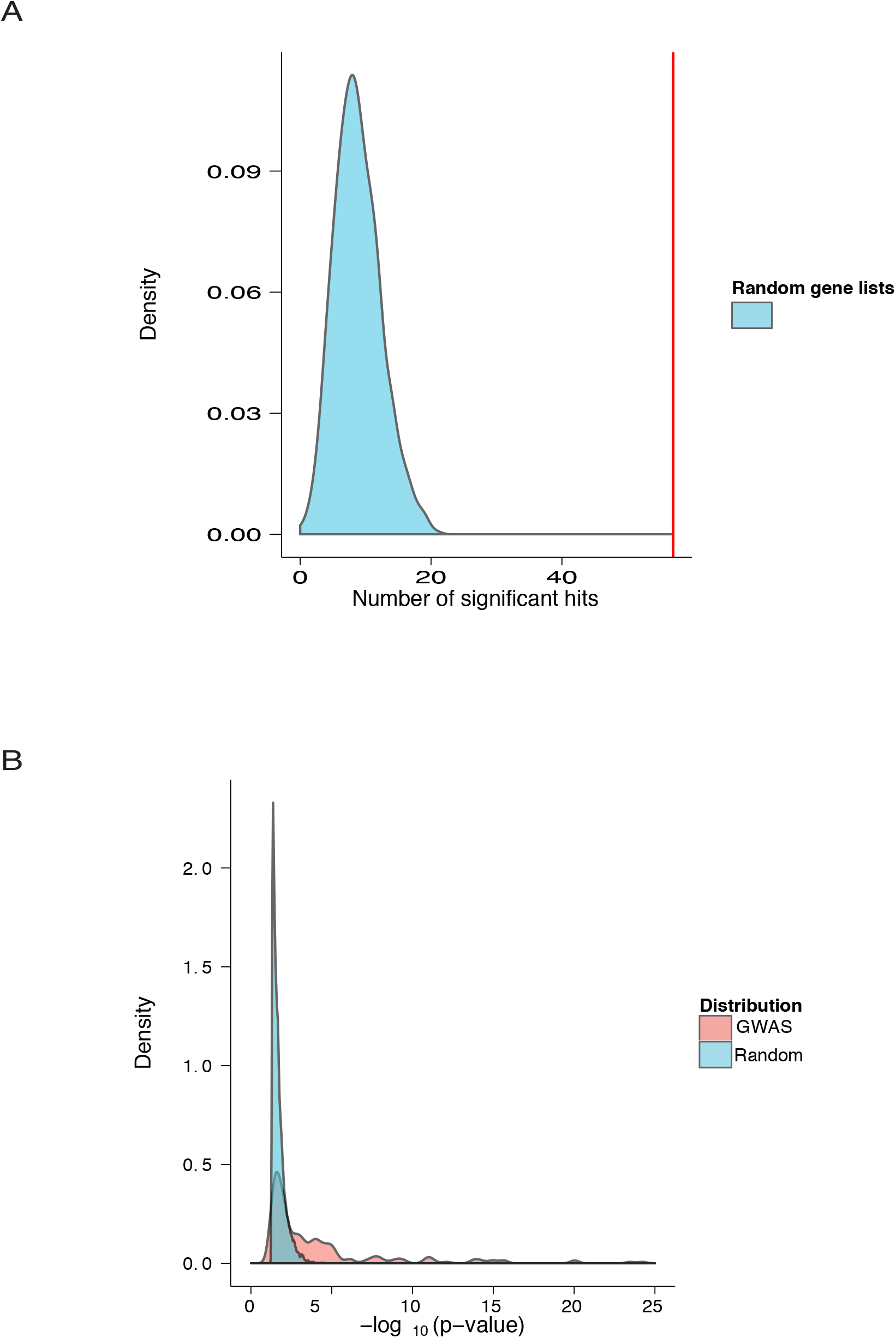
Randomization testing strongly supports the selective expression hypothesis. A) Of the 98 quantitative and disease trait associated gene lists examined, 57 (red line), showed enrichment in at least one tissue. 1000 randomizations of 98 equivalently sized random gene lists resulted in a mean of 8.81 (+/ – 3.53 s.d.) gene lists showing enrichment (blue distribution). B) Median statistical enrichment of those random gene lists showing signal was p-value=0.026 (1.58 as plotted in – log10 scale, blue distribution, range: 0.05 to 2.83E-6), while those from disease and trait associated lists was p-value=0.0027 (2.57, pink distribution, range: 0.05 to 8.50E-28).

To confirm this key result was not a consequence of the manner in which we summarized either the expression data or the GWAS results, we tested the robustness of our analysis to different input parameters. *First,* we were concerned that when a region containing multiple genes was implicated, it is possible that the authors may have been biased in reporting the genes that seemed most plausible to them (i.e. known to be expressed in the relevant tissue). Thus, to avoid this bias we repeated the analysis using the gene-to-SNP mapping reported by the NHGRI staff using a simple, unbiased, proximity-based algorithm. We repeated the analysis with the 87 traits with >30 genes by this mapping strategy and found that there was significant overlap between the candidate gene list and a tissue for 54/87 traits, again a result highly unlikely by chance (Figure S5). Overall, there were 75 traits in common between the 87 traits and the 98 traits described above, and 84% showed concordance between author-mapped and NHGRI- mapped candidate gene lists. The 11 remaining traits typically showed enrichment in a tissue using one mapping strategy but no evidence of overlap with the other mapping strategy, suggesting that the discrepancies were mostly due to false-negatives. Furthermore, out of the 23 traits that only have greater than 30 reported genes when using one mapping strategy but less than 30 in the other, we still see 74% concordance suggesting smaller gene lists can also yield consistent signals. Neither mapping was systematically more sensitive than the other, suggesting author bias contributed little to the signal in the first analysis. *Second*, we confirmed that our results were not strongly biased by the choice of descriptive statistic used for summarizing the expression data. Thus, we also used the median RPKM value of biological replicates as opposed to the mean to make sure that outliers were not artificially inflating expression values of genes, making some genes seem more specifically expressed. When the median is used, 82 out of the 98 (83%) traits highlight similar tissues, and only 3 were discordant. The remaining 13 showed signal with either one summary statistic, or the other, as above suggesting neither approach was systematically more sensitive. *Finally*, we repeated the analysis also using the sub tissue data to confirm that we were not limiting our ability to detect overlaps by averaging across anatomically similar tissues. The results were again highly concordant (71%). Overall, all analyses supported the primary conclusion that disease genes are enriched in relevant tissues, but also suggested that different ways of summarizing data and defining candidate gene lists can capture slightly different relationships, just as running the same gene list through slightly different implementations of GO-based analyses usually highlights similar, but not identical pathways across tools.

Also, note that the challenge of attributing a SNP to a gene is not unique to our work and presents an ongoing problem for all common variant studies. It is a testimony to the robustness of both our method and the finding that the selective expression holds in spite of an unknown level of SNP-to-gene mismapping in the NHGRI summary catalog. This further suggests that our approach may be even more robust and sensitive when applied to contexts where the causative gene is more clear – e.g. exome sequencing studies of *de novo* mutation or sets of genes discovered from mendelian disorders. Indeed, TSEA results for several traits using highly penetrant genes identified using these rare variant methods (Figure S6) identified clear signal in the relevant tissue: genes that had multiple de novo mutations identified in children with autism or epilepsy have signal in the brain ^56,57^. Thus, our approach is readily applicable to results from rare variant analyses as well.

### TSEA indicates skin-expressed loci are key regulators of white blood cell count

We next sought to test whether this now-established relationship between anatomical expression and disease risk might be used to provide insight into the tissues that actually mediate complex traits. One initially unexpected TSEA result was that genes associated with the quantitative trait white blood cell count (WBC) are enriched in the set of transcripts that are found highly expressed in the skin, rather than transcripts from the blood, where the WBC are themselves most abundant (Figure 6A). This human finding joins others from model organisms suggesting that there may be an important pleiotropic relationship between the genetic factors contributing to skin integrity and the regulation of this hemopoeitic trait. The first evidence for this relationship was reported in mice with an allelic series of skin specific knockouts in the Notch pathway or the fatty acid transporter gene *Slc27a4/Fatp4*^11^, which led the authors to suggest that a barrier defect resulted in increased TSLP, which drove WBC. The increase in TSLP and WBC was proportional to the macroscopic assessment of the severity of the skin lesions produced by the different LOF alleles. However, a direct measurement of barrier function, such as transepidermal water loss (TEWL), and its correlation to WBC has not been reported. This leaves open the possibility that the previous WBC increase seen in Notch mutants was caused directly by failed skin differentiation, rather then barrier dysfunction specifically.

**Figure 6.**
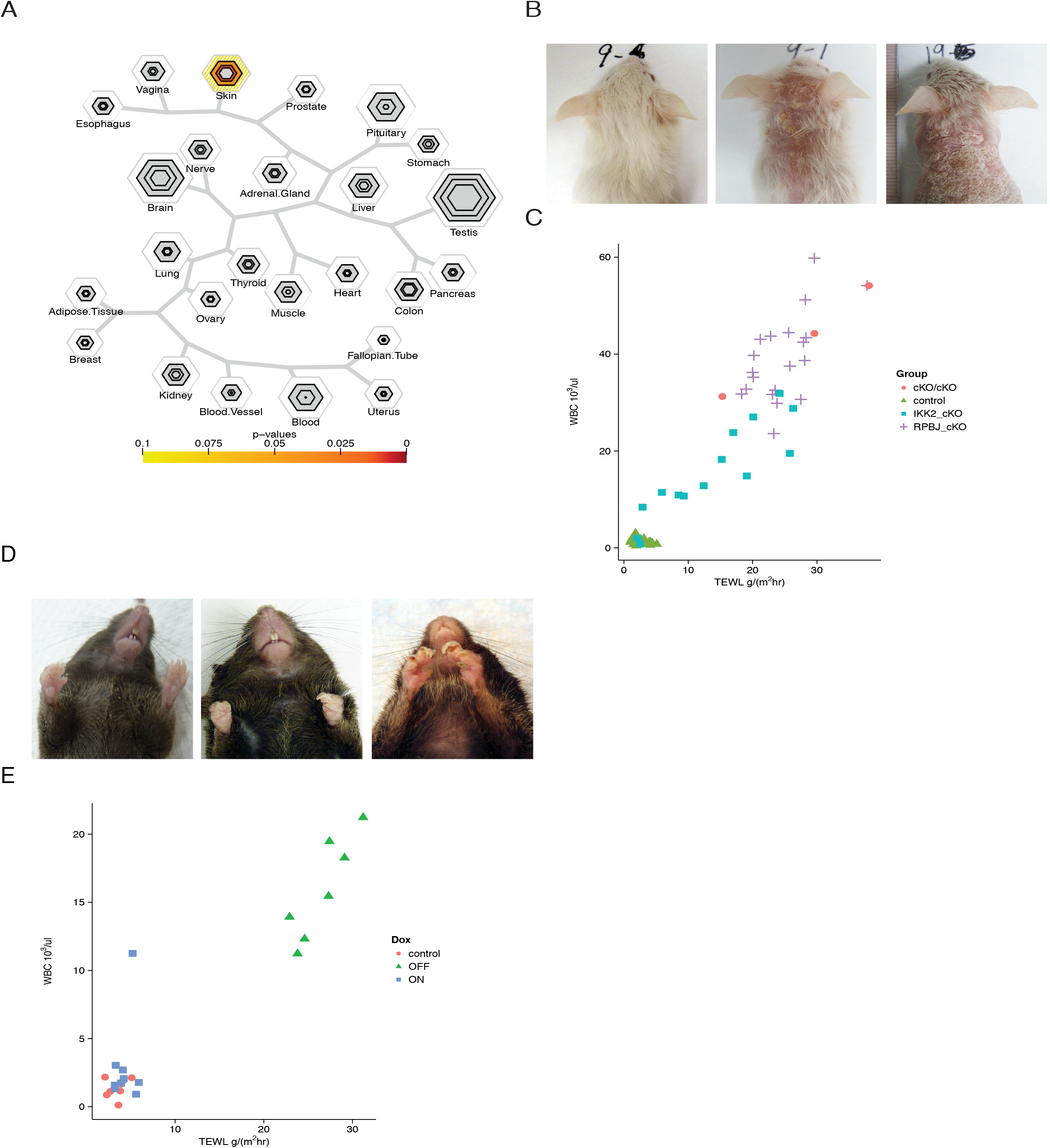
TSEA can identify novel relationships between candidate genes and tissues. A) Genes found to be associated with white blood cell count are disproportionately transcribed in the skin. B) Wild type control mouse, *Ikk2* cKO mouse, and *Rbpj* cKO mouse (from left to right) C) White blood cell count (10^3^cells/ml) as a function of TEWL (g/m^2^hr) in control mice (n=26), *Rbpj* cKO mice (n=20), *Ikk2* cKO mice (n=14), and double cKO *Rbpj/Ikk2* mice (n=3). D) Wild type control mouse, *Lamc2* rKO mouse on Dox, and *Lamc2* rKO mouse off dox E) White blood cell count as a function of TEWL in control mice (n=8), *Lamc2* rKO mice on dox (n=9), *Lamc2* rKO mice off dox (n= 7).

We set out to distinguish these possibilities by taking a quantitative measure of skin integrity, TEWL. In addition, the candidate gene list for human WBC we culled from the GWAS catalog did not include the Notch gene family nor *TSLP* gene, and the TSEA analysis suggested that the relationship between skin expressed genes and WBC exists across many different molecular genetic pathways impacting the skin barrier. Thus, we tested three lines of mutant mice with genetic lesions including two in distinct, Notch-independent pathways. These included conditional knockouts of the gene *Rbpj*^58^, a signal mediator in the Notch pathway previously included in our morphological study^11^, and *Ikk2*, a regulator of NF-kB signaling ^59^. Because these pathways are unrelated, we could separate their common impact on the barrier from their specific pathway-related biology. For each of these genes, we used a Cre-Lox deletion strategy to remove the gene only in keratinocytes located along the dorsal and ventral midline (Msx2-Cre^11^), leaving the genes intact within all other tissues including the immune system. As reported, these mutant mice display clear dermatological abnormalities with the *Rbpj* cKOs having a more severe phenotype than the *Ikk2* cKOs (Figure 6B). In all mice we quantitatively evaluated the integrity of the skin barrier TEWL measurements (inside-out barrier function). In parallel we collected blood and determined the WBC count. In all mice we observed a strong correlation between WBC and the integrity of the inside-out skin barrier (Figure 6C, r^2^=0.9006). In addition, as *Ikk2* mutants are thought to have impaired sensing of outside-in barrier function, the results with these mice suggest that pathogen infiltration (outside-in) barrier function may not be necessary to trigger elevated WBC.

We also tested the quantitative relationship between barrier integrity and WBC using a milder model of skin disruption. Junctional epidermolysis bullosa (JEB) is a skin blistering disease, most often caused by mutations in one of the three chains of laminin (Lm)-332, the α3, β3, or γ2 chains ^60–66^. Mice that lack expression of Lm-332 die shortly after birth with blistering of the skin and oral mucosa ^67–69^. We have rescued the laminin γ2-deficient (*Lamc2^-/-^*) mice by expressing a dox-controllable human laminin γ2 transgene under the keratinocyte-specific K14 promoter (*Lamc2* rKO)^70^. In the absence of dox in their drinking water, these mice gradually develop phenotypes similar to that observed in JEB. Unlike the models above, these mice do not develop severe lesions upon their backs, and undergo normal skin differentiation. Rather, upon withdrawal of doxycycline the mice evidence a macroscopic slight reddening of all skin and the presence of blistering on the paws (Figure 6D), along with additional microscopic evidence of JEB. TEWL and WBC measures were collected from age-matched animals either maintained or withdrawn from doxycycline. Here we show again that as the skin barrier loses integrity, WBC increases (Figure 6E, r^2^=0.8956). These manipulations clearly provide quantitative validation of previous Notch analyses and show that genetic modifiers with a range of molecular mechanisms for decreasing skin integrity consistently increase WBC. This supports the suggestion that polymorphism which regulates skin genes in humans may also contribute to phenotypic variation in human WBC.

## Discussion

To build a foundational resource for expression based ‘pathway-like’ analyses, we have systematically defined sets of genes with enriched expression across a range of tissues, and tested the utility of this resource for several applications. First, we have formally tested the long-held assumption that diseases genes overall should be enriched in their expression in the tissues afflicted by the disease. Second, we defined the tissues implicated by this assumption for every trait available in the NHGRI catalog (Movie S1). Third, we provided a clear additional proof-of-principle analysis in mice for the insights that might be provided by this approach with independent biological experiments demonstrating a quantitative relationship between the genetics of skin integrity and the immunological trait of white blood cell count. Finally, we show that our resource and approach have applications to gene sets derived from data sources beyond GWAS. It is also worth noting that the TSEA method is not limited to disease gene lists and can be used to highlight tissues related to gene lists from any analysis. For example, using the recently identified set of the 1003 most constrained genes in the human genome^56^, we observe a robust overrepresentation of brain, nerve, and pituitary expression (Figure 7). This finding is noteworthy because it suggests that much of the evolutionary pressures of human selection might be occurring on genes mediating behavior, rather than other physiological traits.

**Figure 7.**
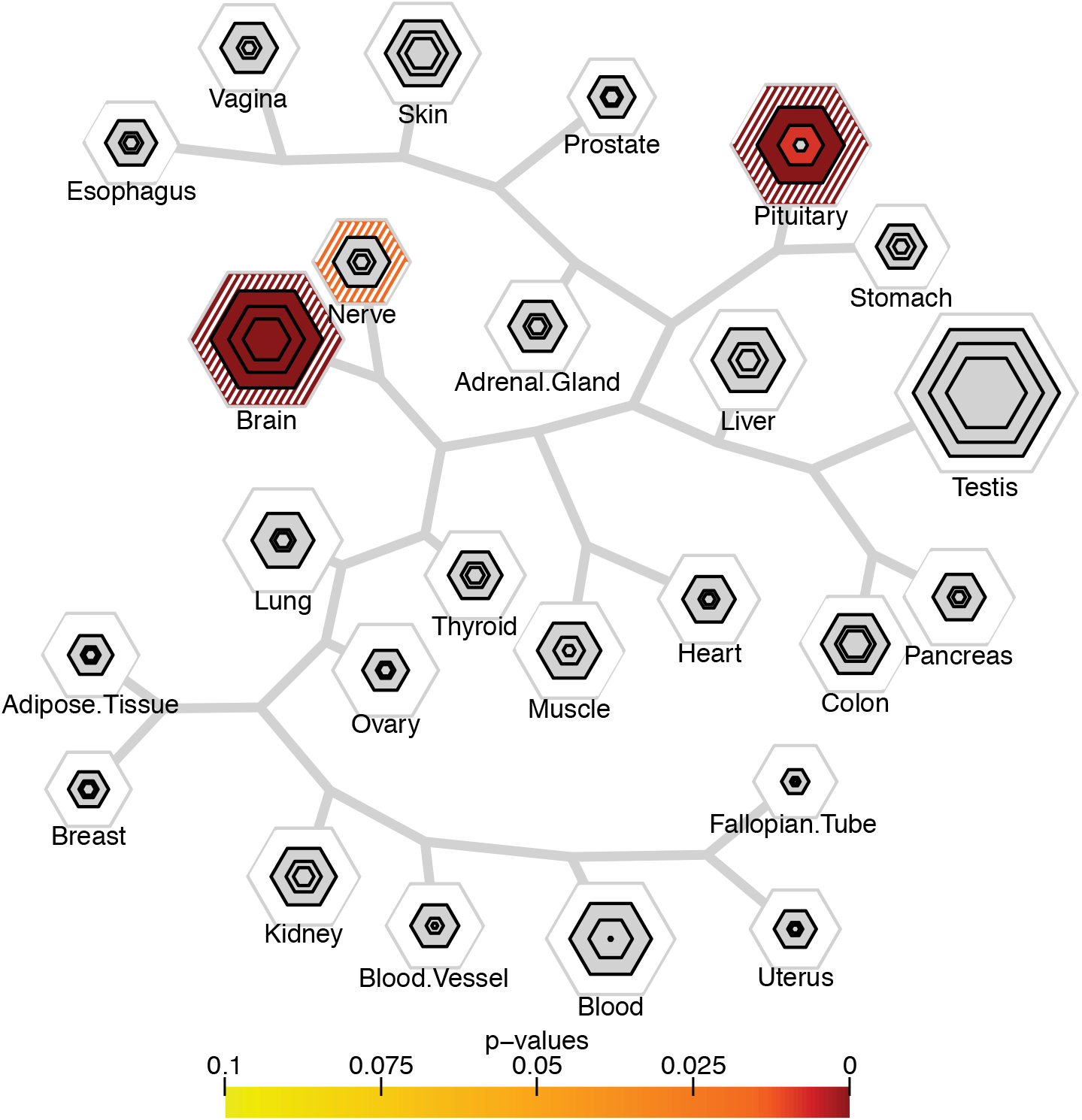
Constrained genes are expressed in the brain, nerve, and pituitary. The top 1003 constrained genes as identified by^56^ show an over representation of genes expressed in the brain, pituitary, and nerve.

Note that our conclusions regarding the selective expression hypothesis do not state that every disease gene will follow this pattern, as there are well known examples of broadly expressed genes that still disrupt very specific cell types or tissues (e.g. mutations in the mitochondrial protein SOD1, expressed in every cell, can lead to the specific destruction of motor neurons in Amyotrophic Lateral Sclerosis) ^71^. Rather, the conclusion is that across a large number genes for a given diseases, there will be a relative overabundance of those with enriched expression in the relevant tissue.

Overall, we found 57/98 traits had a relationship to particular tissues. This is a remarkably high proportion, especially given that many of these studies utilized a fairly simple method (genomic proximity) of attributing SNPs to genes, and thus there was likely some added noise in the analysis. We don’t believe this noise influences our main conclusions because previous modeling adding spurious genes actually decreased our power to detect tissue-specific enrichment^9^, and thus would contribute only to false negatives rather than false positives. Nonetheless, false negatives here should be interpreted with caution until future analyses incorporating new data better integrating gene expression and genomic polymorphisms are included. These, along with results from exome sequencing studies, will likely improve our ability to ascribe genetic risk to particular genes and thus improve the sensitivity of the method.

The overall support for the selective expression hypothesis presented here has at least three clear practical future applications. 1) In situations where the genetic analyses are robust but the biological mechanisms are not well understood, TSEA may be used to identify the relevant tissues. For example, GWAS signals for uric acid levels clearly identify the kidney as the source of the metabolite (Figure 3). For future disease biomarkers of unknown tissue source, GWAS in conjunction with TSEA may provide some biological insight. 2) Systematic use of expression data has the potential to help prioritize variants discovered in GWAS or exome studies. Furthermore, in studies where the relevant tissue is known, TSEA could provide biological prior information for GWA analysis, which could increase the power of studies to detect disease loci, or help to determine which SNPs or genes within large LD blocks might be the most relevant. This is an area that will be pursued in the future. 3) Even when the relevant tissue is thought to be well understood, TSEA may provide for novel insights into the data. For example, we were initially surprised to see data indicating that BMI is regulated by genes expressed in the brain, but in the last three years this has become accepted by human geneticists studying obesity with the tentative explanation that genes regulating appetite probably contribute substantially to weight ^33^. Here, we have provided a clear additional proof-of-principle analysis for this approach for WBC and skin integrity, suggesting the causative alleles in the human QTL studies might also act via modifying skin integrity.

Thus, to facilitate hypothesis generation and other analyses by the scientific community, we have implemented a simple TSEA tool on our website (http://genetics.wustl.edu/jdlab/tsea/). More importantly, we have provided an R package to conduct these analyses (pSI), and provided pre-calculated pSI values for each tissue type (R package on the website pSI.data, or Table S1) to permit other researchers to apply these approaches to their own data, as well as to facilitate the propagation of these sets of tissue enriched genes into other tools for gene set enrichment analysis^2–5^.

## Experimental Procedures

### Ethics statement

All procedures using mice were approved by the Washington University School of Medicine Animal Studies Committee and were performed in accordance with the Animal Welfare Act and the NIH Guide for the Care and Use of Laboratory Animals. The mice were housed in a pathogen-free barrier facility under the veterinary care provided by the Division of Comparative Medicine. Euthanasia was performed using carbon dioxide.

### GTEx dataset

The gene expression data used is the previously normalized set of RNA-Seq data downloaded from the GTEx project (GTEx Analysis 1/31/13 data release, summarized to genes)^6^. This GTEx dataset is comprised of 1,839 samples derived from 189 post-mortem subjects, and more complete description will be available in their forthcoming manuscript. The 1/31/13 release included samples from 45 different tissues, with some tissues offering multiple ‘sub-tissue’ types (i.e., multiple brain dissections). To clearly analyze the data at the tissue level, RPKM values for the sub-tissue types were averaged resulting in 25 ‘whole-tissue’ types. The average number of distinct human samples per sub-tissue type was 36.9 samples and the tissue-level aggregation of the data yielded an average of 66.4 samples per tissue, and all but two had n>6. Thus, the transcripts detected as enriched in each tissue here should be fairly representative of the population, with the possible exceptions of the kidneys (n=3), and fallopian tubes (n=1) which should be regarded more tenuously. To prepare the already normalized data for input to the pSI algorithm, biological replicates were averaged and genes were filtered to include well-annotated protein-coding genes designated by RefSeq (release 60) gene annotations. After filtering, 18,056 transcripts remained.

To summarize the variance of sample expression across tissue types, the multivariate total sum of squares (TSS) is calculated as the sum of squared Euclidian distances of individual expression profiles to the average expression across all samples (i.e., centroid of the expression profiles). The within group sum of squares (WSS) is calculated as the sum of squared distances of expression profiles to the corresponding within group average of expressions. The within group variance component is calculated as WSS/TSS. The statistical significance if this measure is estimated by permutation.

### pSI values and enrichment analysis

Using the pSI R package function specificity.index ^7,9^, the RPKM values for each transcript were used to calculate a pSI value for that transcript in each of the tissues. pSI values were calculated for all detectably expressed (>0.3 RPKM, as per ^13^) protein coding transcripts. As an independent method of generating tissue specific gene lists, we calculated the Shannon entropy as described^14^ and used a similar permutation method that is implemented in the specificity.index function to assign p-values to the Shannon entropy measures. Correlations between the SI and entropy values of genes were assessed using the Spearman correlation (Figure S3).

To analyze the enrichment of a candidate gene list in tissues, we calculated the significance of the overlap between the candidate gene list and the transcripts enriched in each tissue using the Fisher’s Exact test. P-values were further adjusted using a Benjamini-Hochberg (B-H) correction across tissues to account for multiple comparisons.

### RPKM heatmaps

To evaluate the expression of genes specific to tissues at various thresholds, the RPKM values for those genes were visualized using the R package gplots. At each pSI threshold (0.05, 0.01, 0.001, 0.0001), the union of all tissue-specific genes was taken and their respective log-transformed RPKM values were plotted as heatmaps. For visualization, samples were hierarchically clustered at the least stringent pSI threshold (0.05) and the resulting order was kept for all figures. To prevent infinite values after log-transformation, genes with RPKM values equal to zero had a constant of 1E^-5^ added. Similarly, the overlap of the gene list for each tissue at each threshold was plotted as a heatmap and hierarchically clustered to observe the amount of list-wise sharing between tissue specific gene lists for the sub-tissue data (Figure S1).

### Validation of TSEA tool with Gene Ontology (GO)

For every tissue in GTEx, we identified a GO term associated with the development of that tissue. All human genes associated with that term were downloaded and subjected to TSEA. ‘On Target’ hits were defined as a significant enrichment in the appropriate tissue. For comparison, random lists of genes, consisting of the same number of genes as those for each GO term, were also selected and analyzed by TSEA.

### Candidate gene lists

The GWAS catalog is a curated dataset of GWA studies with the variants and genes that were reported associated with 940 traits and diseases^10^. We only used traits that had more than 30 associated genes; 98 traits passed this filter. To avoid any bias using the author reported genes, we also performed the TSEA analysis using genes the SNPs mapped to based on proximity, determined by NHGRI staff. When SNPs were reported as intergenic, we used the closest of the up and downstream genes. Previous modeling work indicated that >30 genes provides sufficient power to detect enrichment in these analyses ^9^ The exome gene lists are described previously (ref the epilepsy paper and Daly paper). The top 1003 constrained genes are described in (ref Daly paper).

### Testing the selective expression hypothesis

We first characterized the distribution of results from the true candidate gene lists for each of the 98 GWAS traits to the transcript-enriched list of each tissue from the Fisher’s Exact test, implemented in the fisher.iteration function in the pSI R package^9^. Significant p-values after B-H correction indicate an overrepresentation of a trait’s candidate gene list in at least one tissue’s transcript-enriched list. We compared the number of real traits with at least one tissue identified by TSEA to an empirically derived null distribution of the same number of ‘traits’ with at least one tissue identified by using 1000 sets of random gene-lists size-matched for each GWAS trait (98,000 iterations total). This analysis was repeated using 87 GWAS traits and mapped genes provided by NHGRI.

### Skin integrity mouse models

All procedures using mice were approved by the Washington University School of Medicine Animal Studies Committee and were performed in accordance with the Animal Welfare Act and the NIH Guide for the Care and Use of Laboratory Animals. The mice were housed in a pathogen-free barrier facility under the veterinary care provided by the Division of Comparative Medicine. *Msx2-Cre/+; RBP-j^f/f^* (*RBPj* cKO)^11^ and *Lamc2*^*-/-*70^; *K14-rtTA; TetO-LamC2+* (*Lamc2* rKO) mice have been described previously and were maintained on mixed genetic backgrounds. To generate *Msx2-Cre/+*; *Ikk2^f/f^* (*Ikk2* cKO) mice, *Msx2-Cre/+* transgenic mice ^59,66^were bred with *Ikk2^f/f^* mice ^72^, and the offspring were intercrossed. Doxycycline (dox; 1 mg/ml) was provided in the drinking water of *Lamc2* rKO mice from conception until 8 weeks old, and then omitted from their drinking water for 6 weeks prior to transepidermal water loss (TEWL) and white blood cell (WBC) measurements. *RBP*j cKO and *Ikk2* cKO mice were examined at 3 weeks of age.

TEWL, a marker of epidermal skin barrier function, was measured using a VapoMeter (Delfin Technologies) directly on the *RBPj* cKO and *Ikk2* cKO mice, as these mice exhibit hair loss in the dosal midline region. Hair was removed from *Lamc2* rKO mice using Nair depilatory cream 24 hr prior to examination. Blood samples were collected and the WBC counts were measured using the Hemavet 950 analyzer (Drew Scientific). These studies were performed using separate groups of mice at least three independent times.

## Author Contributions

A.W. performed the pSI analysis and developed the R package. N.K. performed the GWAS analysis and randomization experiments. D.O. created the hexagon plots for graphical display. W.Y. implemented Shannon entropy and explored GTEx data. X.X. and A.N. provided statistical support. T.A-K. and R.K generated the mouse models and collected blood and TEWL measurements. J.D. and D.O. performed GO term validation of TSEA. J.D. supervised experiments. A.W., N.K., and J.D. wrote the manuscript; all authors contributed to the editing of the manuscript.

## Acknowledgements

The authors would like the thank D. Conrad, J. Lowe, I. Borecki, and N. Parikshak for helpful discussions and analytical assistance. The authors declare no conflicts of interest. We would like to thank J.Abu-Amar and M. Pasparakis for mouse lines.

The authors would also like to thank S. Volpi, K. Ardlie, and the researchers of the Genotype-Tissue Expression (GTEx) Project.

